# Microscale Characterization of Undecalcified Bone Tissue Using Correlative Atomic Force Microscopy and Histology

**DOI:** 10.64898/2026.10.16.759704

**Authors:** Nada Wahman, Young Hun Kim, Jahkwaun Budhai, Bikhyat Shrestha, Francesca Mazzoni, Crystal Huang, Ning Fan, Eric Rosiek, Eric Chan, Murray J Tipping

## Abstract

**SUMMARY:** Characterizing bone tissue in its mineralized state remains technically challenging. Conventional approaches typically compromise mechanical integrity through decalcification or lack the structural resolution necessary to capture microscale heterogeneity. Here we present an integrated workflow combining atomic force microscopy (AFM) with histological and immunofluorescence assays on adjacent undecalcified bone tissue cryosections. Using a tape-transfer mounting approach, we resolve mechanically and structurally distinct tissue types within a minimally processed single mouse femur section. Furthermore, we show that decalcification substantially alters nanoscale surface topography and local mechanical properties. Applied to an aging cohort of C57BL/6J female mice spanning 7 to 115 weeks, the proposed method detects age-associated changes in bone matrix structure and stiffness. The approach provides a practical methodology for spatially correlating nanomechanical properties with molecular markers in bone and other mineralized tissues, with broad applicability to disease models including cancer metastasis, osteoporosis, and osteoarthritis.

## INTRODUCTION

Bone is a hierarchically organized composite material whose structural integrity arises from interactions between mineralized collagen fibrils and a complex cellular network ^1,2^. The multiscale architecture allows bone to withstand mechanical loading and resist fracture ^3,4^. Characterization of mineralized bone tissue is essential for understanding skeletal health, disease progression, and fracture susceptibility. However, conventional mechanical assays, such as bulk compression and three-point bending, provide useful measurements of overall bone mechanics, but primarily capture macroscopic properties and obscure heterogeneity within the matrix ^4–6^. Structural imaging methods spanning the micro- to nanoscale, such as micro-computed tomography (micro-CT) and scanning electron microscopy (SEM), have improved structural assessment of bone; however, important limitations remain. Micro-CT primarily reconstructs mineralized architecture without direct assessment of local material properties ^7^. Alternatively, SEM requires dehydration and conductive coating that can alter tissue structure and prevent quantitative mechanical measurements ^8^.

Atomic force microscopy (AFM) addresses many of these constraints by enabling quantitative nanoscale mapping of surface topography and local Young’s modulus as a measure of stiffness in minimally processed specimens ^9–11^. In mineralized tissues, AFM nanomechanical analysis has revealed spatial heterogeneity in Young’s modulus across the mineralized matrix at sub-micron resolution ^12,13^. Topographic imaging has complemented these mechanical measurements by visualizing collagen fibril organization and mineral tessellation patterns directly from the bone surface ^14–17^. AFM has also been applied to investigate how the mechanical properties of cells and their environment influence metastatic behavior and cancer progression in breast cancer bone metastasis ^18^ and melanoma bone metastasis ^19^ models. Despite these advances, nanomechanical measurements alone lack the biological context needed to fully interpret localized changes in tissue function, particularly in the setting of aging ^20–22^ or disease, where multiple compositional changes co-occur. Correlative approaches combining molecular and mechanical information have been established in other connective tissues, such as immunofluorescence-guided AFM mapping of the pericellular matrix mechanics in cartilage ^23,24^, yet their application to mineralized bone tissue remains limited.

While histological and immunofluorescence methods provide highly specific molecular information, they are traditionally conducted on decalcified tissue, limiting their ability to preserve and assess structural features alongside molecular targets. Decalcification precludes comprehensive evaluation of the mineral phase, a major contributor to bone stiffness, and, depending on the protocol, can perturb collagen structure and molecular integrity ^25–27^. Consequently, decalcification can limit the ability of samples to fully reflect the compositional and mechanical environment of bone. Methods for sectioning undecalcified bone exist ^28^, including the Kawamoto tape transfer cryosectioning technique ^29^. However, their use for spatial correlation of AFM measurements with histological and immunofluorescence data from adjacent sections remains limited. Extending such correlative approaches to emerging spatial-biology platforms, including transcriptomic profiling ^30–32^, could provide molecular resolution beyond immunofluorescence and enable more comprehensive spatial characterization of the bone microenvironment.

Here we propose an integrated method combining AFM with histological and immunofluorescence assays on adjacent undecalcified mouse femur cryosections from the same tissue block. The approach enables spatial correlation among biochemical composition, local stiffness, and nanoscale topography of the bone microenvironment. We first demonstrate that the workflow resolves mechanically distinct tissues within a single mouse femur section. We then compare undecalcified and EDTA-decalcified bone to quantify the effects of mineral removal on bone material properties. Finally, we apply the workflow to a murine aging cohort spanning skeletal maturation to senescence, demonstrating the method’s sensitivity to progressive biological change. By linking matrix composition to local mechanical properties, the proposed methodology provides a comprehensive workflow for functional characterization of mineralized bone tissue.

## RESULTS

### A tape transfer workflow enables correlative AFM and histological analysis of undecalcified bone sections

To enable joint nanomechanical and histological bone profiling in its mineralized state, we developed an integrated sample preparation and analysis workflow (Figure 1). Femora from C57BL/6J female mice (Figure 1a, left) were embedded, fresh frozen, and cryosectioned without prior decalcification, maintaining the bone’s mineral and collagen composite architecture. Frozen samples were equilibrated in a cryostat (Figure 1a, I.) and sectioned using a tungsten carbide blade (Figure 1a, IV.) equipped with the Kawamoto tape transfer system (Figure 1a, middle) ^29^. Adhesive tape was applied to the trimmed block prior to sectioning (Figure 1a, II., III.), supporting the undecalcified section during cutting. The approach allowed for a clean transfer onto the glass slides without fracture or curling (Figure 1a right), overcoming a common challenge when sectioning mineralized tissue. Sections were cut at 10 µm for AFM analysis to ensure sufficient material for force indentation measurements and at 5 µm for histological staining to facilitate optimal staining penetration and image clarity. Mounted sections were fixed with 4% paraformaldehyde in PBS and kept on ice until analysis ^33^.

**Figure 1.**
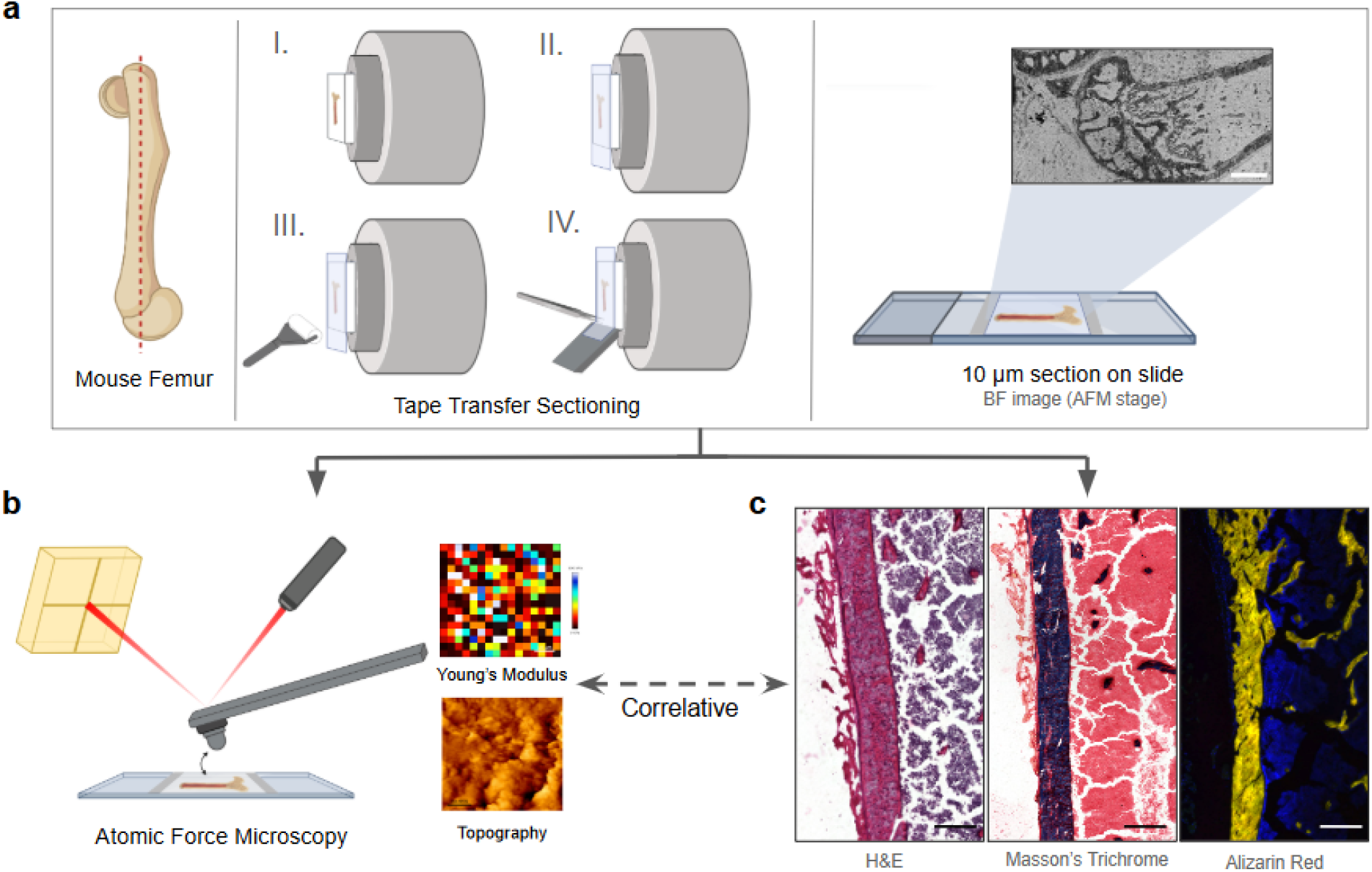
Workflow for correlative AFM, histological, and fluorescence analysis of undecalcified bone tissue. **(a)** Fresh-frozen mouse femora (left) are cryosectioned using the Kawamoto tape transfer method (I-IV), and sections are mounted on glass slides. Representative brightfield image of a 10 µm section on the AFM stage (right). Scale bar, 500 µm. **(b)** Schematic of AFM characterization and representative Young’s modulus and topography maps. The dashed double-headed arrow indicates correlative analysis with adjacent tissue sections. **(c)** Representative H&E, Masson’s Trichrome, and Alizarin Red fluorescence imaging of adjacent sections. Scale bar, 250 µm.

Adjacent sections designated for AFM were first imaged using brightfield microscopy on the AFM stage to generate an overview map for region-of-interest selection (Figure 1a, right). AFM characterization yielded independently acquired topography and Young’s modulus maps, providing complementary nanoscale readouts of surface topography and local mechanical properties (Figure 1b). Sections from the same specimen were processed for histological and immunofluorescence staining, enabling spatial correlation of nanomechanical properties and structural markers from corresponding tissue regions (Figure 1b-c). A summary of all analytical modalities employed in this workflow, including their targets, underlying principles, and the information they provide, is presented in Table 1.

**Table 1.** Summary of complementary characterization modalities employed in this workflow.

| Modality | Target | Principle | Information provided |
| --- | --- | --- | --- |
| Hematoxylin and eosin (H&E) | Nuclei, cytoplasm, and extracellular matrix | Hematoxylin binds to negatively charged nucleic acids (blue/purple signal); eosin binds to positively charged proteins in the cytoplasm and extracellular matrix (pink signal) | General tissue morphology and cellular architecture |
| Masson's Trichrome | Connective tissue and cellular components | Aniline blue stains collagen; Biebrich scarlet stains muscle and cytoplasm | Delineation of connective tissue compartments, including collagen, muscle, and bone marrow |
| Alizarin Red | Calcium deposits | Alizarin red S forms a chelate complex with calcium ions, (red/orange signal) | Distribution and abundance of minerals within the bone matrix |
| Collagen I immunofluorescence | Type I Collagen protein | Primary antibody specifically binds the target protein; fluorescent secondary antibody enables visualization | Spatial distribution and relative expression of the primary structural matrix protein |
| Osteopontin immunofluorescence | Osteopontin protein | Primary antibody specifically binds the target protein; fluorescent secondary antibody enables visualization | Spatial distribution and relative expression of a protein involved in bone mineralization and mineral-matrix interactions |
| AFM topography | Surface architecture | Cantilever deflection maps surface height variations at nanoscale resolution via QI mode (JPK NanoWizard V) | Nanoscale surface features reflecting mineral and collagen organization |
| AFM force mapping | Local Young's modulus | Force-distance curves yield Young's modulus via the Hertz contact mechanics model using Contact Mode Force Mapping (JPK NanoWizard V) | Local tissue stiffness quantified by Young's modulus |

*AFM, atomic force microscopy; H&E, hematoxylin and eosin; QI, quantitative imaging.*

### AFM and histological analysis of a single bone section resolve mechanically and structurally distinct tissue types

The workflow’s performance was assessed across multiple tissue types present within a single minimally processed mouse femur section to demonstrate that AFM can resolve microscale heterogeneity across compositionally distinct materials. Longitudinal sections of the murine femur encompass bone, skeletal muscle, and bone marrow, which span several orders of magnitude in mechanical stiffness with molecular profiles distinguishable by histology ^12,25^. Representative adjacent sections were stained with hematoxylin and eosin (H&E) for general tissue morphology and Masson’s Trichrome (Table 1), which distinguishes collagen (blue) and muscle (red). Sections were mapped via AFM in parallel to spatially register nanomechanical properties onto the underlying tissue structure (Figure 2).

**Figure 2.**
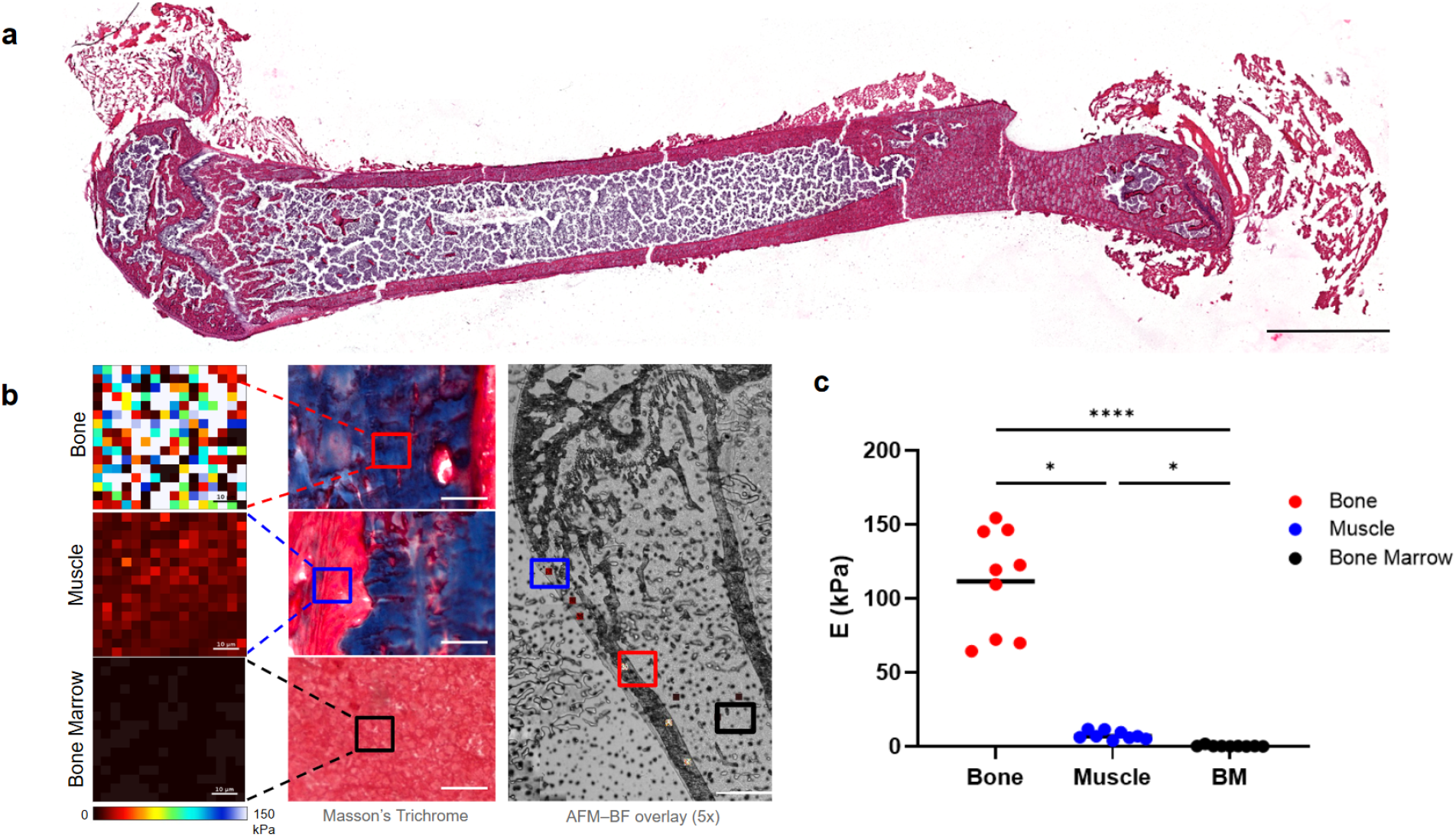
Correlative AFM and histological characterization of bone, muscle, and bone marrow. **(a)** H&E-stained whole femur 5 µm section from a 7-week-old mouse. Scale bar, 2.5 mm. **(b)** Representative Young’s modulus maps (0–150 kPa) of bone, muscle, and bone marrow (left). Scale bars, 10 µm. Masson’s Trichrome staining of corresponding tissue regions (center). Scale bars, 30 µm. AFM Young’s modulus maps overlaid onto a 5x brightfield image of the tissue section (right). Scale bar, 500 µm. **(c)** Young’s modulus *(E)* across bone, muscle, and bone marrow. Each point represents an individual force map, and horizontal lines indicate the median. Statistical significance was determined using a Kruskal–Wallis test followed by Dunn’s multiple-comparisons test; \**p* < 0.05, \*\*\*\**p* < 0.0001. *n* = 9 force maps per tissue type from three sections from three mice.

A whole-section H&E overview, representative of standard histological staining, demonstrates that the workflow preserves tissue morphology and provides a guide for region-of-interest selection during AFM acquisitions (Figure 2a). Masson’s Trichrome staining of adjacent sections reveals delineation of bone, bone marrow, and surrounding muscle, confirming compatibility with connective tissue stains (Figure 2b center). The histologically defined tissue regions were compared against AFM-derived mechanical boundaries (Figure 2b left), with AFM force maps overlaid onto brightfield images acquired with a 5x objective (Figure 2b right). Young’s modulus measurements confirmed that bone exhibited the highest stiffness, followed by muscle and bone marrow (Figure 2c). Bone stiffness was significantly greater than both muscle and bone marrow (p = 0.0485 and p < 0.0001, respectively), and muscle was significantly stiffer than marrow (p = 0.0485), consistent with the established mechanical hierarchy of these tissues ^12^. Force mapping was additionally performed using stiffer AFM probes to access the higher stiffness regime of mineralized bone tissues, yielding modulus values in closer agreement with previously reported values for mineralized bone ^14,34^ (Figure S1c,f). Furthermore, Young’s modulus maps acquired at false-color scales appropriate to each tissue type indicate mechanical heterogeneity within individual compartments (Figure S2a). Together, the approach resolves distinct mechanical properties across tissue types within a mineralized mouse femur section while remaining fully compatible with subsequent histological analysis of adjacent sections.

### Characterization of nanomechanical properties and surface morphology in decalcified and undecalcified bone

Decalcification is the chemical removal of the mineral phase from bone tissue, typically achieved using chelating agents such as ethylenediaminetetraacetic acid (EDTA) ^26^. The process is routinely employed in bone research to facilitate sectioning and immunostaining, as it renders mineralized tissue compatible with standard histological workflows ^28^. Prior studies have demonstrated that demineralization alters the bulk mechanical properties of cortical bone at the tissue level ^6,27^. Microscale imaging techniques, including scanning electron microscopy, have demonstrated decalcification alters collagen fibril organization and bone surface morphology ^35,36^. However, the consequences of decalcification on nanoscale surface morphology and local mechanical properties as characterized by AFM remain incompletely characterized. To examine these effects, sections of femora from 7-week-old mice were first processed in the undecalcified state to preserve mineral and collagen composite structure. Contralateral femora from the same mice were processed using EDTA-based decalcification for comparison ^26^ prior to sectioning. Both conditions were subjected to identical AFM topography and force mapping protocols (Figure 3). The side-by-side characterization aimed to clarify what information each preparation method preserves or sacrifices at the nanoscale to support more informed decisions about which approach best suits a given research question.

**Figure 3.**
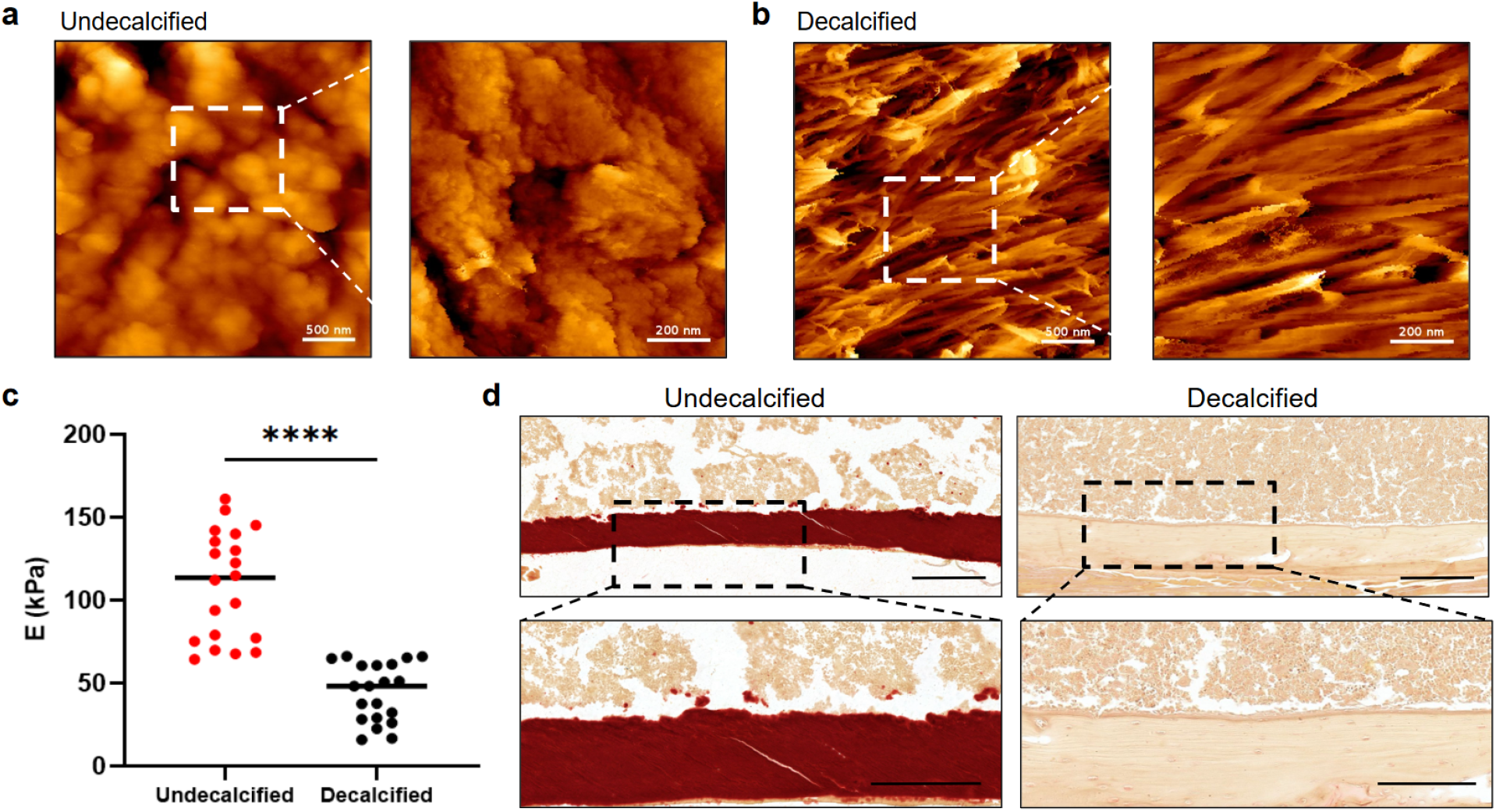
Nanoscale structural and mechanical characterization of undecalcified and decalcified bone. **(a–b)** Representative AFM topography images of undecalcified **(a)** and decalcified **(b)** bone from 7-week-old mice. Images were acquired at 3 × 3 µm (scale bars, 500 nm), with corresponding 1 × 1 µm higher-resolution images (scale bars, 200 nm). **(c)** Young’s modulus *(E)* of undecalcified and decalcified bone. Each point represents an individual force map, and horizontal lines indicate the median. Statistical significance was determined using a two-tailed Mann–Whitney test; \*\*\*\**p* < 0.0001. *n* = 20 force maps from three sections from two mice per condition. **(d)** Alizarin Red staining of adjacent 5 µm sections of undecalcified (left) and decalcified (right) bone from 7-week-old mice. Overview images (top) and corresponding enlarged regions (bottom) are shown. Scale bars, 500 µm (overview) and 250 µm (enlarged regions).

AFM topography of undecalcified 7-week-old bone revealed a heterogeneous, granular surface, characteristic of the mineralized collagen fibril composite ^17^ (Figure 3a). High-resolution 1 µm x 1 µm scans (Figure 3a right, white dashed box), acquired within larger 3 µm regions (Figure 3a left), revealed discrete surface features with height variations on the order of tens of nanometers. In contrast, decalcified sections exhibited a transition to elongated, aligned fibrillar features, consistent with uncovering of the underlying collagen fiber network following mineral removal ^36,37^ (Figure 3b). Each preparation thus captured a distinct aspect of bone morphology. The morphological differences were accompanied by a significant decrease in Young’s modulus. Undecalcified bone exhibited a broad distribution of modulus values ranging from 70-150 kPa, indicative of mechanical heterogeneity of the mineralized state (Figure 3c). Decalcified sections exhibited a significantly lower and narrower distribution, with values predominantly below 100 kPa (Figure 3c; p < 0.0001), consistent with loss of the mineral phase and its contribution to bone stiffness.

Alizarin Red, a dye that forms a chelate complex with calcium ions producing an intense red signal (Table 1), was applied to adjacent sections to assess mineral content differences underlying these nanomechanical observations (Figure 3d) ^38^. Undecalcified sections displayed deep crimson staining throughout the bone matrix (Figure 3d left), indicating retained calcium content. In contrast, the decalcified bone sections appeared pale (Figure 3d right), indicating reduced calcium content. The observation is further supported by Alizarin Red staining with Light Green counterstaining of adjacent sections (Figure S3a and S3b), which provided contrast for visualization of non-mineralized tissue structures and was consistent with substantial removal of the mineral phase during EDTA treatment. Taken together, the AFM and histological data provide a multimodal, correlative characterization showing the impacts of decalcification on nanoscale surface morphology and local mechanical properties. The findings offer researchers a quantitative reference for understanding the tradeoffs between preparation methods when designing studies of bone microenvironment mechanics and composition.

### Age-dependent decline in bone stiffness and compositional changes detected by multimodal analysis

The methodology was employed on bone sections from C57BL/6J female mice across a comprehensive cohort spanning 7 to 115 weeks of age to demonstrate the biological applicability of the workflow (Figure 4) ^39,40^. Aging was selected as a model because it encompasses the full transition from skeletal growth to senescence to illustrate the method’s sensitivity to progressive biological change. Female mice were used to reduce sex-related variability in bone composition and because age-related bone density loss is more pronounced in females than in male C57BL/6J mice ^39^. AFM topography scans of femoral bone from young (7-week) and aged (100-week) mice revealed qualitative differences in surface topography (Figure 4a). Femoral bone from young mice displayed pronounced granular structures and clearly defined surface features (Figure 4a top) compared to the smoother, more uniform appearance of aged bone (Figure 4a bottom). Height profile traces taken along the white arrowed lines in the scans confirmed higher variation in young bone compared to the lower, more uniform profiles of aged femurs (Figure 4a right). The observations suggest structural changes to the bone surface with advancing age (Figure 4a; Figure S2b).

**Figure 4.**
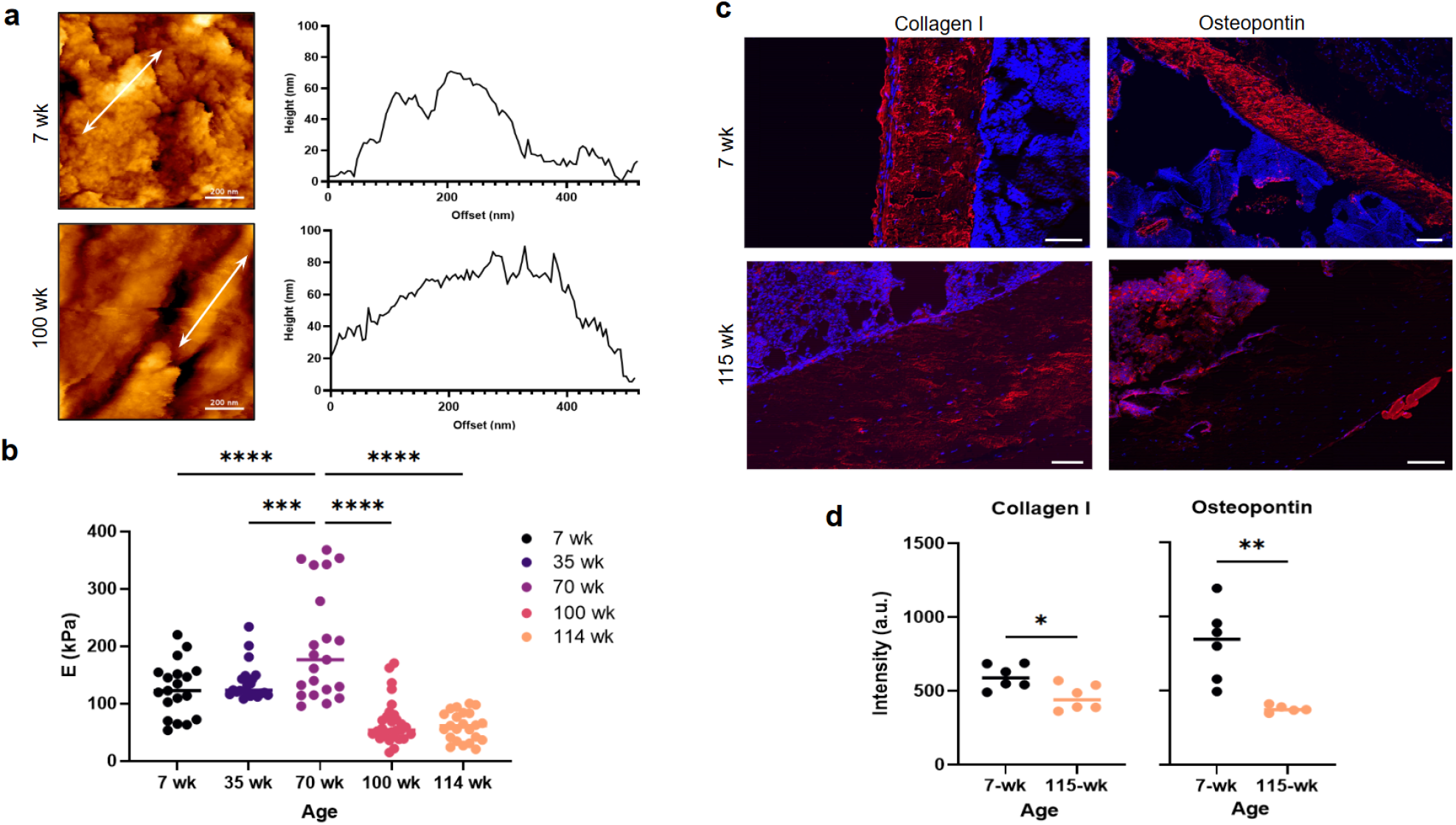
Structural, mechanical, and molecular characterization of bone across age. **(a)** Representative AFM topography and corresponding height profiles of bone from 7-week-old (top) and 100-week-old (bottom) mice. White arrows indicate the 500 nm paths used to generate the corresponding height profiles. Scale bars, 200 nm. **(b)** Young’s modulus *(E)* across five age groups. Each point represents an individual force map, and horizontal lines indicate the mean. Statistical significance was determined using an ordinary one-way ANOVA followed by Dunnett’s multiple-comparisons test, with the 70-week age group used as the comparison group; \*\*\**p* < 0.001, \*\*\*\**p* < 0.0001. *n* = 19–29 force maps per age group from 4–5 sections from 1–3 mice per age group. **(c)** Representative Collagen I (left; red) and Osteopontin (right; red) immunofluorescence images of bone from 7-week-old (top) and 115-week-old (bottom) mice, with DAPI shown in blue. Scale bars, 50 µm for Collagen I and 100 µm for Osteopontin. **(d)** Quantification of Collagen I and Osteopontin fluorescence intensity in 7- and 115-week-old bone. Horizontal lines indicate the mean. Collagen I fluorescence intensity decreased by 23.6% and Osteopontin fluorescence intensity decreased by 53.8% in 115-week-old compared with 7-week-old bone. Statistical significance was determined using two-tailed unpaired *t*-tests with Welch’s correction; \**p* < 0.05, \*\**p* < 0.01. *n* = 1 section from one mouse per age group and condition.

Young’s modulus mapping across five age groups revealed a biphasic trajectory, outlined by an initial increase during skeletal maturation followed by a progressive decline with advanced age (Figure 4b). Stiffness increased from the 7-week age group (black) and remained elevated at 35 weeks (dark purple), reaching its highest values at 70 weeks (purple). Notably, the 70-week age group displayed a bimodal distribution of Young’s modulus values (Figure 4b). Stiffness then declined sharply in the older 100-week (pink) and 114-week (peach) age groups (Figure 4b). Young’s modulus at 70 weeks was significantly greater than at 7 weeks (p < 0.0001), 35 weeks (p = 0.0004), 100 weeks (p < 0.0001), and 114 weeks (p < 0.0001). The increased data dispersion of modulus values at 70 weeks suggests greater local mechanical heterogeneity within the cortical matrix (Figure 4b) ^20,22^.

Complementing these age-dependent mechanical and structural changes, adjacent sections from 7-week and 115-week mice were immunostained for collagen I and osteopontin (Figure 4c). Immunofluorescence staining enabled spatial visualization of protein expression within tissue sections by using antibodies that bind target proteins with high affinity, with signal intensity reflecting relative protein abundance ^41^. Collagen I, the primary structural protein of the bone matrix, provides the scaffold upon which minerals are deposited and is a key determinant of tissue architecture (Figure 4c left) ^1,2^. Collagen I immunofluorescence signal intensity was reduced in the aged femoral bone compared with young (23.6% decrease, p = 0.0167), accompanying the smoother surface topography in aged bone AFM scans relative to the pronounced granular features of young bone (Figure 4a, Figure 4d left). Osteopontin, a non-collagenous matrix protein involved in bone mineralization and mineral–matrix interactions, also showed reduced immunofluorescence signal intensity in the aged femur (53.8% decrease, p = 0.0081) (Figure 4c right, Figure 4d right) ^42^. The reduction in osteopontin signal accompanied the lower Young’s modulus detected by AFM in aged mouse femur.

### DISCUSSION

Here we develop a correlative method to combine AFM topographical and nanomechanical characterization with histological and immunofluorescence imaging on adjacent undecalcified murine femoral cryosections. By leveraging the Kawamoto tape transfer technique ^29^, we demonstrate that AFM measurements can be obtained from cryosectioned bone without the need for prior demineralization or extensive processing ^28,34^. The proposed methodology preserves the mineralized bone matrix while maintaining compatibility with histological, immunofluorescence, and spatially resolved molecular analyses. The mechanical properties of bone are closely linked to the molecular and structural composition ^1,2^. Spatial correlation of nanomechanical, molecular, and structural information from adjacent sections therefore offers greater insight into the compositional influence on mechanical properties in undecalcified bone tissue.

Initial validation of the method across mechanically and compositionally distinct tissue types (i.e. bone marrow, muscle, bone) within a single mouse femur established the sensitivity of the AFM measurement approach. Absolute Young’s modulus values for bone measured by AFM vary substantially across the literature and depend heavily on probe stiffness, sample preparation, hydration state, and the analytical model used. Studies using stiff probes on polished or resin-embedded bone sections often report modulus values in the GPa range ^14,34^. In contrast, studies using soft colloidal probes on fresh, minimally processed bone report much lower effective modulus values, reflecting mechanical properties at the scale of the local tissue microenvironment ^12^. The values reported here are specific to the preparation and measurement conditions used. As with all AFM-based mechanical measurements, absolute values should be interpreted relative to the specific probe geometry, cantilever stiffness, and contact model applied, rather than as absolute tissue stiffness values ^43,44^. Measurements performed with stiffer probes (Figure S1e) yielded higher Young’s modulus values, demonstrating the method’s sensitivity across different experimental setups. Importantly, the relative mechanical hierarchy, in which bone is significantly stiffer than both muscle and marrow (p < 0.05 and p < 0.0001, respectively), and muscle is stiffer than marrow (p < 0.05), is consistent with the expected mechanical behavior of these tissues ^12^. Successful H&E and Masson’s Trichrome staining of adjacent tape transfer sections illustrates AFM and histological co-analysis can be performed on adjacent sections from the same block. The approach therefore enabled spatially informed interpretation of nanomechanical data within the broader tissue structure.

The comparison of undecalcified and EDTA-decalcified sections from contralateral femora provides quantitative nanoscale insight into how each preparation method captures distinct aspects of bone structure and mechanics. In undecalcified bone, AFM topography captured the mineralized surface morphology, while force mapping yielded Young’s modulus values spanning 70-150 kPa. Decalcified sections exposed the underlying collagen network through topographical mapping, with a more compressed modulus distribution predominantly below 100 kPa. The observations are consistent with prior reports on the importance of mineral content in governing cortical bone mechanics ^27,34^ and extend this understanding to the nanosurface morphology resolved by AFM topography. Alizarin Red staining further corroborates the differences, supporting an association between the mechanical and structural changes and mineral loss ^38^. Taken together, the choice of preparation method shapes what information is accessible and should be guided by the research question being addressed. Importantly, the workflow described here can be applied to both undecalcified and decalcified preparations. The preparation method can therefore be selected based on the structural, mechanical, or molecular information of interest. Undecalcified and decalcified preparations thus provide distinct and potentially complementary windows into bone biology, and an informed choice between them enables a more comprehensive characterization of the bone microenvironment.

Application of the workflow to an age-dependent cohort of C57BL/6J female mice spanning 7 to 115 weeks demonstrated the method’s ability to detect age-associated differences in molecular and nanomechanical properties. The observed modulus trajectory, with stiffness increasing through skeletal maturation and declining with advanced age, is consistent with known biology of bone aging ^40,45^ and with prior AFM-based reports of age- and tissue-maturation-related changes in bone mechanics ^22,46^. Here, the decline in Young’s modulus in aged mice was accompanied by reduced collagen I and osteopontin immunofluorescence intensity, consistent with the known compositional changes ^20,40^ that occur during aging. Reduction in osteopontin immunofluorescence signal intensity in aged bone suggests decreased protein abundance in the matrix. Given osteopontin’s role in bone mineralization and mineral–matrix interactions ^42^, reduced protein abundance may contribute to the mechanical changes observed by AFM in aged femur.

The bimodal distribution of Young’s modulus values observed in the 70-week age group warrants further consideration. The pattern may reflect increased local mechanical heterogeneity, where tissue regions may differ in mineral density or remodeling activity ^6,20,46^. One potential contributor is the osteocyte lacunar-canalicular network, as the perilacunar matrix has been shown to exhibit distinct mechanical properties relative to the surrounding mineralized tissue that vary with local remodeling activity ^47^. Periosteal and endosteal cortical surfaces have also been shown to differ in mechano-responsiveness and osteocyte density ^48^. The spatial resolution of AFM force mapping is well-suited to capture such local mechanical heterogeneity. Future studies targeting defined anatomical regions, such as periosteal versus endosteal surfaces ^48^ or regions with differing osteocyte lacunar density ^47^, could elucidate the structural basis of this heterogeneity. Smaller AFM tips resolving more localized mechanical contributions may further clarify whether the bimodal distribution reflects spatially discrete regions of bone with differing remodeling or mineralization states.

Beyond these methodological considerations, the approach described here is applicable to other mineralized tissues and disease models. Bone metastasis represents one particularly relevant application given the mechanical properties of the bone microenvironment are increasingly recognized as important regulators of metastatic behavior. Matrix stiffness in the bone microenvironment has been shown to influence cancer cell behavior and metastatic progression ^19,49^, making mechanical characterization of mineralized bone tissue relevant to understanding disease colonization. AFM has already been applied to characterize the mechanical properties of breast cancer bone metastases ^18^, indicating that metastatic lesions exhibit distinct nanoscale mechanical signatures compared to surrounding bone tissue. Age-related changes in bone composition and mechanics may therefore alter the mechanical environment encountered by metastatic cells during bone colonization. The methodology could also be applicable to models of osteoporosis, osteogenesis imperfecta, and fracture healing to investigate alterations in tissue composition, organization, and bone mechanics. More broadly, spatially correlating local nanomechanical properties with molecular markers could provide a framework for investigating mechanoreciprocity across diverse biological and disease contexts. The workflow is therefore adaptable to a wide range of research questions and extends the spatial and mechanical characterization of disease beyond what histological and immunofluorescence staining provides alone.

That said, AFM and immunofluorescence operate at differing spatial scales, and the adjacent section design means that corresponding regions, rather than identical locations, are being compared. As a result, direct causal links between specific molecular changes and local mechanical properties cannot be established from these datasets alone, though the regional correlations observed provide a basis for more targeted future studies. A key advantage of AFM is the ability to vary tip diameter to access different length scales within the same tissue, where larger tips capture tissue-scale mechanical responses while smaller tips provide sensitivity to more localized mechanical features ^50^. Variation in tip diameter therefore offers multi-scale mechanical mapping and, when combined with spatially resolved molecular data from the same specimen, could establish more direct spatial associations between local protein expression and nanomechanical behavior. Furthermore, super-resolution fluorescence microscopy has been demonstrated on clinical tissue sections and fresh frozen cryosections ^51,52^. Such approaches could be applied to sections generated by this workflow, enabling immunofluorescence imaging at spatial scales to be more closely matched to localized AFM measurements and improving spatial correlation between molecular and mechanical features.

The ability to analyze undecalcified adjacent cryosections provides opportunities to integrate molecular approaches beyond histological and immunofluorescence staining. For example, spatial -omics platforms can offer transcriptomic data at single-cell resolution for researchers seeking deeper insights into cell expression and cell state. The data can elucidate the basis of structural and mechanical changes observed by AFM, extending beyond what conventional staining alone can provide ^53,54^. As molecular profiling protocols for mineralized tissues continue to advance ^54,55^, adjacent sections could in principle be designated for AFM force mapping, immunofluorescence, and spatial -omics, enabling complementary mechanical and molecular readouts from matched tissue regions. Together, these extensions could enable increasingly comprehensive integration of nanomechanical and molecular information within the bone microenvironment.

In conclusion, we demonstrate an integrated workflow that combines AFM with histological and immunofluorescence analysis on adjacent undecalcified bone cryosections. The method preserves the mineralized bone matrix while enabling spatial comparison of local mechanical properties and tissue composition. Using this approach, we resolved mechanically distinct tissue compartments, characterized the structural and mechanical consequences of decalcification, and detected age-associated changes in bone mechanics and composition. Together, the findings establish a practical methodology for investigating the molecular and structural components of bone mechanics in aging, disease, and other mineralized tissue models.

## METHODS

### Bone tissue collection and preparation

Femora from female C57BL/6J mice were provided by the Tuomas Tammela Laboratory at Memorial Sloan Kettering Cancer Center (MSKCC) and the Elaine Fuchs Laboratory at The Rockefeller University. All animal studies conducted at MSKCC were approved by the MSKCC Institutional Animal Care and Use Committee (IACUC; protocol number 17-11-008), and MSKCC guidelines for the proper and humane use of animals in biomedical research were followed. Animals at The Rockefeller University were maintained and bred under specific-pathogen-free conditions at the Comparative Bioscience Center (CBC), an Association for Assessment and Accreditation of Laboratory Animal Care (AAALAC)-accredited facility.

Following collection, femora were embedded in optimal cutting temperature (OCT) compound (Fisher Scientific) for subsequent cryosectioning. Samples were frozen in small baths of isopentane on dry ice and stored at −80°C until sectioning. Samples from mice ranging from 7 to 115 weeks of age were used for the aging cohort.

### Cryosectioning and Tape Transfer

Bone samples were cryosectioned using a Leica CM1950 cryostat. The cryostat chamber temperature was maintained between −20°C and −30°C and the head of the cryostat around −20°C to −25°C. Samples were placed in the cryostat and allowed to equilibrate to the temperature for approximately 30 minutes before sectioning.

Samples were sectioned using the 40-UT tungsten carbide blade (Section-Lab) and the angle of the blade holder was set to match the manufacturer’s guidelines for the 40-UT blade. The blocks were mounted on a chuck with OCT and trimmed until a desired surface of the tissue was exposed. Kawamoto adhesive tape (Cryofilm type 3C(16UF), Section-Lab Co.Ltd.) was cut to a desired width and placed in the cryostat to cool. Prior to sectioning, the surface of the trimmed block was warmed gently with a finger, and the adhesive side of the Kawamoto tape was then placed onto the block surface. A roller was used to firmly adhere the tape to the tissue surface and remove air bubbles. While holding the non-adhesive edge of the tape with forceps, the samples were sectioned evenly at a thickness of 10 µm for AFM analysis and 5 µm for histological and fluorescence analysis.

Following sectioning, the Kawamoto tape containing the tissue sections was mounted onto a labeled glass slide with the tissue facing upward. For sections designated for AFM analysis, double-sided tape was placed beneath the Kawamoto tape to secure it to the slide. Sections designated for histological and fluorescence analysis were secured to the slide at the corners using autoclave tape.

AFM sections were maintained on ice with aluminum foil placed beneath the slide and fixed for 10 min using cold 4% paraformaldehyde (PFA) in PBS. Sections were then washed three times with cold PBS and maintained on ice in PBS until AFM measurements were performed on the same day.

For decalcification experiments, bone samples were fixed overnight in 4% PFA in PBS at room temperature, washed with water, and decalcified in 0.5 M EDTA (pH 8) for approximately 5 days. Following decalcification, samples were washed, post-fixed overnight in 4% PFA in PBS at room temperature, and processed for OCT embedding and cryosectioning.

### Atomic Force Microscopy

#### Force Mapping

AFM measurements were conducted with a NanoWizard V (JPK-Bruker) in Contact Mode Force Mapping at room temperature in PBS. For tissue force mapping, a 10 µm diameter spherical AFM probe (nominal spring constant k = 0.2 N/m, SAA-SPH-5UM, Bruker) was used. Each AFM probe’s spring constant was measured by the thermal noise method. A setpoint of 1 nN and an approach velocity of 15 µm/s were used (60 µm x 60 µm image size with 16 × 16-pixel resolution). The setpoint was selected to maintain indentation within approximately 10 – 20% of the section thickness and minimize influence from the underlying glass substrate. Force curves from each force map were fitted using the Hertz contact mechanics model to determine Young’s modulus:

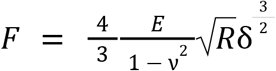

Where *F* is the applied force, *E* is the Young’s modulus, ν is Poisson’s ratio, *R* is the probe radius, and δ is the indentation depth. A Poisson’s ratio of 0.5 was assumed for the tissue. Data was processed with JPK Data Processing software.

To measure the higher stiffness range of mineralized bone tissue, force mapping was additionally performed using stiffer 10 µm diameter colloidal probes (nominal spring constants k = 5.3 N/m and k = 15.5 N/m, PT.BORO.SI.10.CAL, Novascan) under identical image size and pixel resolution conditions. The setpoint was adjusted for each probe to maintain appropriate indentation depth, as detailed in Figure S1.

#### Topography Imaging

AFM images were captured with a NanoWizard V (JPK-Bruker) in QI™ Advanced mode at room temperature in PBS. For tissue topography imaging, an AFM probe (PEAKFORCE-HIRS-F-A, Bruker) with a nominal frequency of approximately 165 kHz and nominal spring constant of 0.35 N/m was used. Images were collected with a 1 nN setpoint, at image sizes of 3 × 3 µm and 1 × 1 µm with a 256 × 256 pixel resolution. Images were processed using JPK Data Processing software.

### Histological Staining

#### Hematoxylin and Eosin Staining

Sections were dipped in distilled water to remove residual OCT and incubated in Harris Hematoxylin Acidified (6765003, Epredia) for 1 min. Sections were briefly washed in distilled water, incubated in 1x PBS for 1 min, washed again in distilled water, and then incubated in eosin (3801600, Leica) for 1 min. Sections were subsequently dehydrated through two changes of 95% ethanol for 1 min each followed by two changes of 100% ethanol for 1 min each. Slides were cleared in two changes of Sub-X xylene substitute (3803670, Leica) for 1 min each and manually mounted using Permount Toluene Solution mounting medium (SP15-100, Fisher Scientific).

#### Masson’s Trichrome Staining

Sections were incubated overnight in Bouin’s Fluid (159901-01, Electron Microscopy Sciences) and subsequently washed under running tap water. Staining was performed using a Trichrome Stain Kit (ab150686, Abcam). Sections were incubated in Biebrich Scarlet/Acid Fuchsin solution for 1 min, rinsed with distilled water, and incubated for 10 min in a 1:1:2 mixture of phosphomolybdic acid solution (HT153-250ML, Sigma-Aldrich), phosphotungstic acid solution (HT152-250ML, Sigma-Aldrich), and distilled water. Without rinsing, sections were incubated in Aniline Blue solution from the Trichrome Stain Kit for 1 min and then rinsed with distilled water. Sections were dehydrated through two changes of 95% ethanol for 1 min each followed by two changes of 100% ethanol for 1 min each. Slides were cleared in two changes of Sub-X for 1 min each and manually mounted using Permount mounting medium.

#### Alizarin Red Staining

Prior to staining, sections were fixed in 4% PFA in 1x PBS for 10 min and washed three times with PBS. Alizarin Red solution (VB-3008, VitroVivo Biotech) was applied directly to the tissue section and incubated for 1 min. Excess Alizarin Red solution was removed without rinsing, and Light Green counterstain from GMS Stain Kit (ab287884, Abcam) was applied for 1 min. Slides were subsequently dipped 20 times in acetone, followed by 20 additional dips in acetone-xylene solution (1:1). The slides were then cleared in two changes of xylene for 5 min each and manually mounted using Permount Toluene Solution mounting medium.

### Fluorescence and Immunofluorescence Staining

#### Alizarin Red Fluorescent Staining

A staining solution was prepared by dissolving 2 g of Alizarin Red S (A5533-25G, Sigma-Aldrich) in 100 mL of 1% potassium hydroxide. Sections were incubated in the staining solution for 5 min, briefly rinsed in distilled water, and incubated with 5 μg/mL4′,6-diamidino-2-phenylindole (DAPI; Sigma-Aldrich) for 10 min. Sections were then briefly rinsed in distilled water and mounted using Mowiol 4–88 mounting medium (Calbiochem).

#### Immunofluorescence Staining

Immunofluorescence staining was performed on 5 µm fresh-frozen undecalcified mouse femur cryosections prepared using the Kawamoto tape-transfer method. Sections supported on adhesive film were secured to Superfrost glass slides using autoclave tape and processed using an automated Leica Bond RX staining system. Sections were pretreated with EDTA-based epitope retrieval solution (Bond ER2, AR9640, Leica) for 20 min at 95°C.

Primary antibodies against Collagen I (PA1-85319, Invitrogen; 2 µg/mL), Osteopontin (AF808, R&D Systems; 0.25 µg/mL), and CD31 (ab182981, Abcam; 0.08 µg/mL) were applied individually to separate tissue sections and incubated for 1 h at room temperature. Rabbit IgG (10500C, Invitrogen; 3 µg/mL) and goat IgG (005-000-003, Jackson ImmunoResearch; 1 µg/mL) were used as isotype controls for Collagen I and CD31, and for Osteopontin, respectively.Following primary antibody incubation, HRP-based secondary antibody detection was performed using the Leica Bond staining system at room temperature for 8 min. Fluorescent detection was then performed for 10 min using Alexa Fluor tyramide signal amplification reagent (B40953, Life Technologies) or CF dye tyramide 594 conjugate (92172, Biotium).

Following automated staining, slides were washed in PBS and incubated with 5 µg/mL 4′,6-diamidino-2-phenylindole (DAPI; Sigma-Aldrich) in PBS for 5 min and rinsed in PBS. Sections supported on adhesive film were gently repositioned on the glass slides after removal of the autoclave tape. Mowiol 4–88 mounting medium (Calbiochem) was applied, and sections were coverslipped to seal and flatten the tissue. Slides were stored overnight at −20°C before imaging.

### Image Acquisition and Analysis

Histological and fluorescence images were acquired using the Panoramic Scanner (3DHistech) with a 20x/0.8 NA objective. Brightfield images used for correlation with AFM force maps were acquired using a 5x objective. Immunofluorescence images were analyzed using ImageJ. Six regions of interest (ROIs) corresponding to bone tissue were randomly selected per tissue section, and fixed intensity thresholds were applied to quantify the total positive fluorescence signal intensity for collagen I and osteopontin. The resulting fluorescence intensities were used to compare relative fluorescence intensity between young and aged bone.

### Statistical Analysis

Statistical analyses were performed using GraphPad Prism (GraphPad Software). Comparisons of Young’s modulus measurements among bone, muscle, and bone marrow were performed using a Kruskal–Wallis test followed by Dunn’s multiple-comparisons test. Young’s modulus measurements between undecalcified and decalcified bone were compared using a two-tailed Mann–Whitney test. Young’s modulus measurements across the five age groups were analyzed using an ordinary one-way ANOVA followed by Dunnett’s multiple-comparisons test, using the 70-week age group as the comparison group. Collagen I and osteopontin immunofluorescence intensities between young and aged bone were analyzed using two-tailed unpaired *t*-tests with Welch’s correction. Statistical significance was defined as p < 0.05.

## ACKNOWLEDGMENTS

We thank Xueqian Zhuang and the Tuomas Tammela laboratory at Memorial Sloan Kettering Cancer Center, and Jorge Almagro Santiago and the Elaine Fuchs laboratory at The Rockefeller University, for providing mouse tissue samples used in this study. We would like to thank Yevgeniy Romin from the Molecular Cytology Core Facility. This research was supported in part through the NIH/NCI Cancer Center Support Grant P30 CA008748.

## AUTHOR CONTRIBUTIONS

N.W., Y.H.K. and M.T. conceived the project and designed the experiments. N.W. and Y.H.K. developed the AFM methodology and performed the AFM experiments. B.S. contributed to protocol development for bone cryosectioning and histological staining. J.B. performed bone cryosectioning and histological staining and developed the Alizarin Red staining protocol. F.M. developed the protocol and performed the immunofluorescence staining. C.H. contributed to bone cryosectioning and histological staining. N.F. provided histological expertise and guidance. E.C. developed the ImageJ script used for image analysis. E.R. performed slide scanning and optimized image acquisition. N.W. and Y.H.K. analyzed the AFM data. Y.H.K. led the project, with N.W. and Y.H.K. coordinating experimental work across the participating groups. N.W. prepared the figures and wrote the manuscript with input from Y.H.K. and M.T. All authors contributed to discussions and substantively revised the manuscript.

