## Supplemental Information for "Microscale Characterization of Undecalcified Bone Tissue Using Correlative Atomic Force Microscopy and Histology"

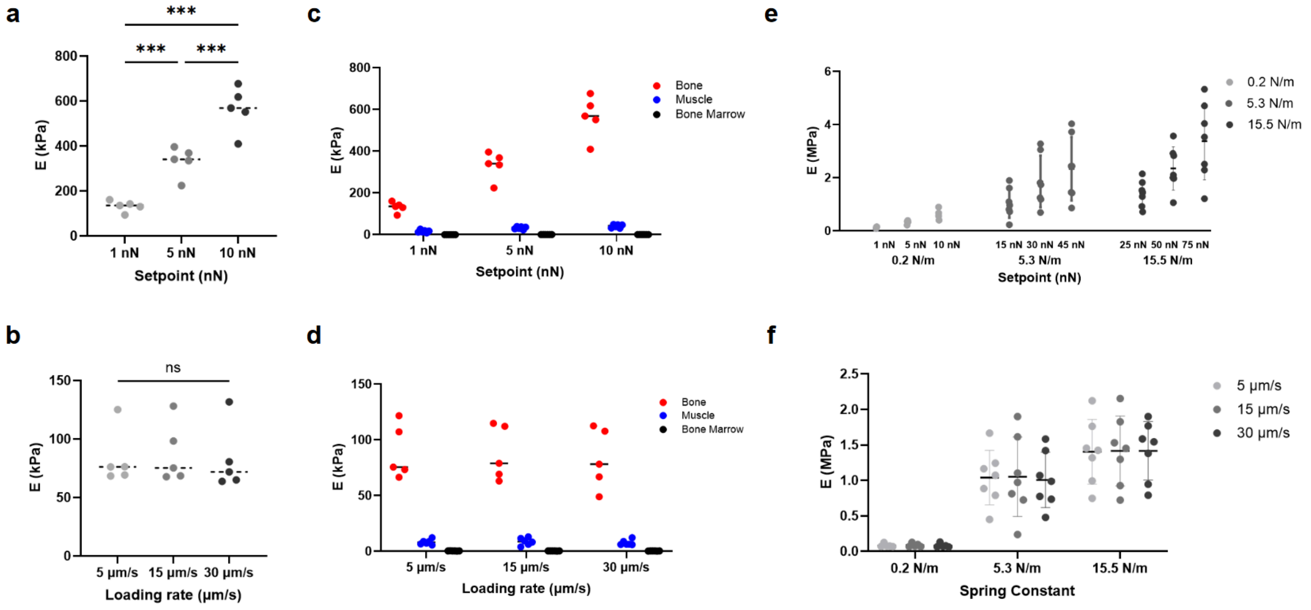


**Supplementary Figure 1. Technical validation of AFM force mapping in tissue sections. (a)** Setpoint dependence of Young's modulus measurements in bone using a 0.2 N/m probe. *$p < 0.05$, **$p < 0.01$, ***$p < 0.001$; repeated-measures one-way ANOVA with Tukey's multiple-comparisons test. **(b)** Loading-rate dependence of Young's modulus measurements in bone using a 0.2 N/m probe. ns, adjusted $p > 0.05$; repeated-measures one-way ANOVA with Tukey's multiple-comparisons test. **(c)** Setpoint dependence of Young's modulus measurements across bone, muscle, and bone marrow using a 0.2 N/m probe. Adjusted *p*-values for setpoint comparisons were $p < 0.006$ for bone and $p < 0.03$ for muscle. For bone marrow, comparisons were $p < 0.003$, except for the 5 vs. 10 nN comparison ($p$ = 0.0604). Two-way repeated-measures ANOVA with Geisser–Greenhouse correction followed by Tukey's multiple-comparisons test. **(d)** Loading-rate dependence of Young's modulus measurements across bone, muscle, and bone marrow using a 0.2 N/m probe. ns, adjusted $p > 0.05$. Two-way repeated-measures ANOVA with Geisser–Greenhouse correction followed by Tukey's multiple-comparisons test. **(e)** Setpoint dependence of Young's modulus measurements using probes with spring constants of 0.2, 5.3, and 15.5 N/m. Adjusted *p*-values for setpoint comparisons were $p < 0.02$ for the 0.2 N/m probe, $p < 0.007$ for the 5.3 N/m probe, and $p < 0.02$ for the 15.5 N/m probe. Repeated-measures one-way ANOVA with Tukey's multiple-comparisons test was performed separately for each probe. **(f)** Loading-rate dependence of Young's modulus measurements using probes with spring constants of 0.2, 5.3, and 15.5 N/m. ns, adjusted $p > 0.05$. Two-way repeated-measures ANOVA with Geisser–Greenhouse correction followed by Tukey's multiple-comparisons test.

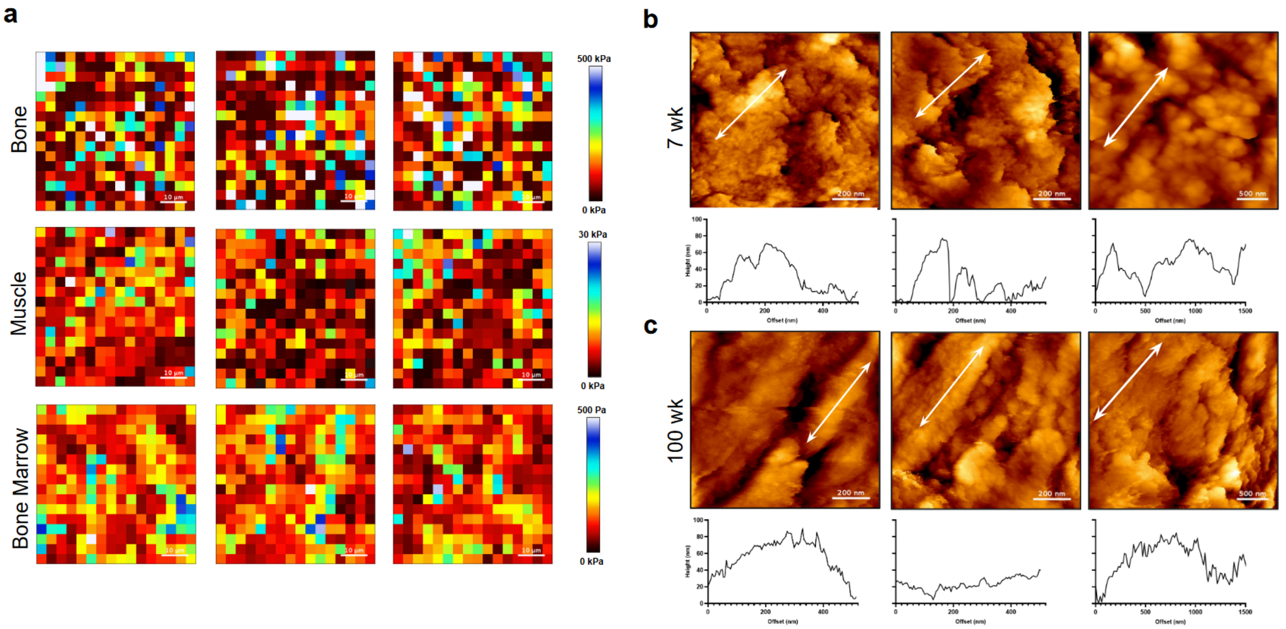


**Supplementary Figure 2. Additional mechanical and topographical characterization of bone tissue. (a)** Representative Young's modulus maps of bone, muscle, and bone marrow. Scale bars, 10 μm. Color scales are independently adjusted for each tissue type to visualize local mechanical heterogeneity. **(b)** Representative AFM topography images and corresponding height profiles from 7-week-old bone. **(c)** Representative AFM topography images and corresponding height profiles from 100-week-old bone. In (b) and (c), the first two columns show 1 x 1 μm scans (scale bars, 200 nm), and the third column shows 3 x 3 μm scans (scale bars, 500 nm). White arrows indicate the locations and directions of the corresponding height profiles, spanning 500 nm for the 1 x 1 μm scans and 1.5 μm for the 3 x 3 μm scans.

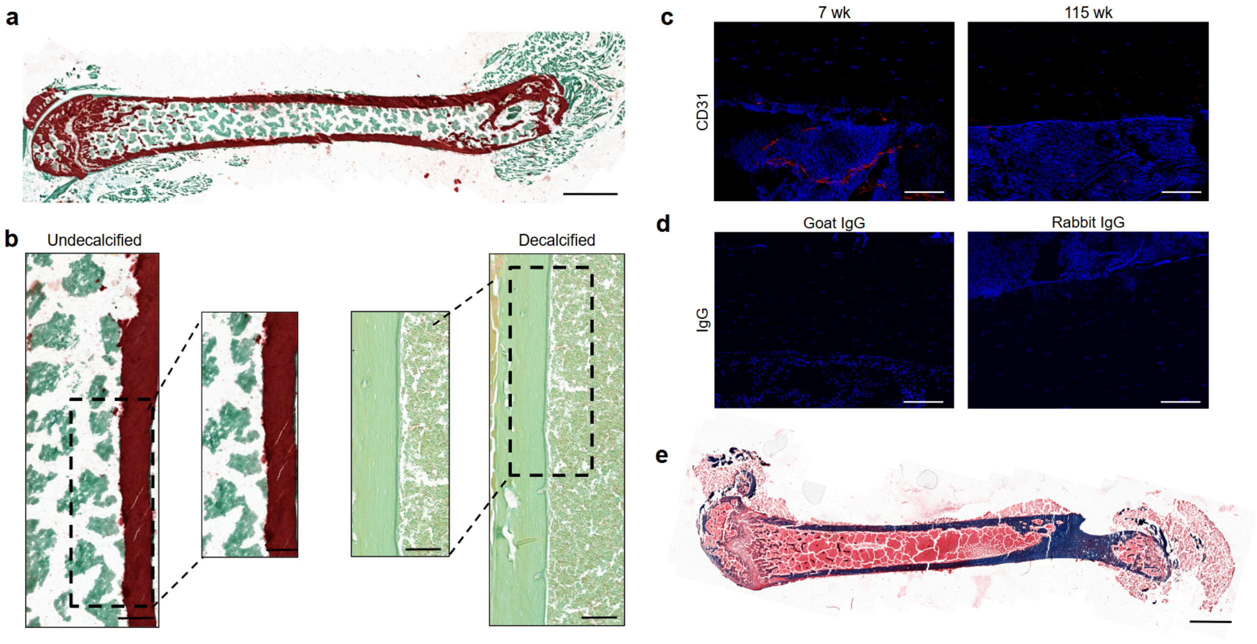


**Supplementary Figure 3. Additional histological and immunofluorescence characterization of bone tissue. (a)** Representative whole-femur image of undecalcified bone from a 7-week-old mouse stained with Alizarin Red and Light Green counterstain. Scale bar, 1 mm. **(b)** Representative enlarged regions of undecalcified and decalcified bone from 7-week-old mice stained with Alizarin Red and Light Green counterstain. Scale bars, 250 µm and 200 µm for undecalcified bone and 125 µm and 100 µm for decalcified bone. **(c)** Representative CD31 (red) immunofluorescence images of bone from 7-week-old (left) and 115-week-old (right) mice, with DAPI shown in blue. Scale bars, 100 µm. **(d)** Representative rabbit and goat IgG isotype controls in bone sections from 7-week-old mice. Rabbit IgG was used as the isotype control for Collagen I and CD31 staining, and goat IgG was used as the isotype control for Osteopontin staining, with DAPI shown in blue. Scale bars, 100 µm. **(e)** Representative whole-femur section from a 7-week-old mouse stained with Masson's Trichrome. Scale bar, 2.5 mm.
